# Detection for endogenous retroviral loci in Jungle fowl using whole-genome sequencing

**DOI:** 10.1101/2023.01.24.525461

**Authors:** Shinya Ishihara

## Abstract

Chickens harbor two primary families of endogenous retroviruses (ERVs): the avian leukosis virus (ALV) and endogenous avian retrovirus (EAV) families. Characterization of ERVs can provide crucial insights into avian evolution. This study aimed to identify novel ERV loci absent in the reference genome using whole-genome sequencing data of red junglefowl, gray junglefowl, Ceylon junglefowl, and green junglefowl. A total of 837 ERV loci were identified across the four Gallus species. The number of ERV loci detected in red junglefowl with its subspecies, gray junglefowl, Ceylon junglefowl, and green junglefowl, was 363, 217, 194, and 129 loci, respectively. The phylogenetic tree was congruent with previously reported trees, suggesting the potential for inferring relationships among past junglefowl populations from the identified ERV loci. Of the detected loci, 306 ERVs were identified near or within the genes, and some were associated with cell adhesion. The detected ERV sequences was in the long terminal repeat region and were classified as EAV-family, ALV-E, Ovex-1, and murine leukemia virus related ERVs. In addition, the sequence of the EAV family was divided into four patterns by combining the U3, R, and U5 regions. These findings contribute to a more comprehensive understanding of the characteristics of chicken ERVs.

## Introduction

Upon retroviral infection, the viral genome is reverse transcribed and integrated into the host genome as a provirus. In principle, the provirus has all the requirements for its replication and consists of an internal region encoding viral genes (gag, pro/pol, and env), which are flanked by two identical regulatory long terminal repeats (LTRs) at integration. Adjacent to the provirus is a short target site duplication (TSD) of 4–8 bp in the host genome sequence generated during integration. Vertical transmission can cause such viruses to infect germ cells and reproductive tissues, resulting in the formation of endogenous retroviruses (ERVs) within offspring. Gradually, ERVs can reach a high frequency within populations and eventually become fixed within species [1]. Typical avian ERVs include the avian leukosis virus (ALV) and endogenous avian retrovirus (EAV) families. The ALV family comprises several subgroups, and the ERVs of subgroup E, referred to as ALV-E, often retain high structural integrity [2]. A sequence known as EAV-HP within the EAV family lacks the pol gene, whereas EAV-0 and EAV-51 have the pol gene but lack the env gene [3]. Chickens and various avian species have been observed to possess gene deletions, suggesting that the EAV family is an ERV that emerges more frequently than ALV-E.

Approximately 5% of the human genome is derived from ERVs, whereas ERVs constitute approximately 3% of the chicken genome [4, 5]. However, there likely exists a significant number of ERVs have not been discovered in chickens. These ERVs have played a role in shaping bird species diversity and have caused economic losses to the poultry industry due to genetic diseases [6, 7, 8]. Characterization of ERVs will provide essential insights into avian evolution.

Mitochondrial DNA analysis indicates that the red jungle fowl is an ancestral species of chickens [9, 10]. In addition to red junglefowl (*G. gallus*), three other species belonging to the genus Gallus were identified: gray junglefowl (*G. sonneratii*), Ceylon junglefowl (*G. lafayetii*), and green junglefowl (*G. varius*). Red junglefowl is distributed across much of Southeast Asia and parts of South Asia, while the other three species have more restricted ranges: gray junglefowl in central and southern India, Ceylon junglefowl in Sri Lanka, and green junglefowl in Java and surrounding islands. Recent molecular genetic studies suggest that various species of Gallus contribute to the genetic composition of fowl. However, the origin and history of genetic diversity in chickens remains only partially understood [11, 12, 13]. In this study, the author aimed to identify ERV loci in the genome using whole-genome data for the genus Gallus, including subspecies. Additionally, by comparing ERV loci among species and detected sequences of ERVs, the characteristics of ERV from the genus Gallus were clarified.

## Result

### Sequencing data quality and identification of the non-reference ERV breakpoint

100 bp paired-end reads were mapped using BWA-MEM [14], and the overall mean sequence depth was 30.6X (13.5–42.9) for all junglefowls (Table S1). The mapping results are presented in Table S1. More than 95.3% of the paired-end reads for each junglefowl were mapped to the reference gallus genome, while only 1.56–31.59% was not properly mapped (improper reads). In addition, 0.10–1.86% of the reads were singletons that were mapped to only one side. The analytical process was conducted in accordance with the methodology of previous studies [15, 16]. Initially, the author employed RetroSeq software [17] for the analysis and identified singleton and improper pairs among the read pairs supporting ERV during the RetroSeq “discover” step. Subsequently, the author utilized the RetroSeq “call” stage results to determine ERV insertion loci (breakpoints). The candidate breakpoint was determined with the filter level adjusted to seven or eight, which is the range used by RetroSeq. The total number of candidate ERV insertion loci identified for each individual ranged from 39 to 2,011 (Table S1). Next, the Integrated Genome Viewer (IGV) [18] was employed to confirm the presence or absence of TSDs for all detected loci on each individual. Furthermore, reads around these loci with confirmed TSD were extracted from the merged bams of each species. Contigs were constructed using the extracted reads and analyzed using blastn [19]. In total, 837 ERV loci were identified. Most of the identified ERVs were related to the LTR region of the EAV family (EAV-HP, EAV-51, EAV-0, ev/J, or chicken endogenous LTR). Twenty LTR sequences of ALV-E were identified, and these sequences were present only in red junglefowl and its subspecies (Table S2). In chr2:133314053, all species and subspecies had a contig similar to the LTR of the murine leukemia virus (MLV) related endogenous retrovirus (DQ280312). Moreover, on chr3:54480182, all species and subspecies, excluding the green junglefowl, had an Ovex1 (FJ406461). Of the ERVs detected, 306 were present near or within the gene (Table S2). The Gene ontology (GO) analysis using these Gene sets showed six GO terms (Table S3). The top-ranked GO category was “cell adhesion” and included genes such as RELN, CNTN5, CDH20, CDH7, TENM1, SPON1, NRXN3, and CDH4.

### Comparison of detected ERV loci among species and subspecies

The number of ERV loci detected in merged red junglefowl, gray junglefowl, Ceylon junglefowl, and green junglefowl was 363, 217, 194, and 129 loci, respectively (Table 1). The number of ERV loci detected in the red junglefowl and its subspecies ranged from to 61–124. The Venn diagram shows ERVs with shared loci among subspecies or species (Figure 1A and 1B). Among the species, 53 loci were detected in common between two or more species, with gray junglefowl and Ceylon junglefowl exhibiting the highest degree of commonality among species, at 37 loci in common. In contrast, no common loci were detected between green junglefowl and red junglefowl. Among the subspecies of red junglefowl, 13 common ERV loci were detected, with the most common loci identified between the red junglefowl and G.spadiceus (Thai), at 58 shared ERVs. The clustering tree created based on all the loci is shown in Figure 1C. The tree branched in the following order: green junglefowl, Ceylon junglefowl, gray junglefowl, and red junglefowl.

**Figure 1.**
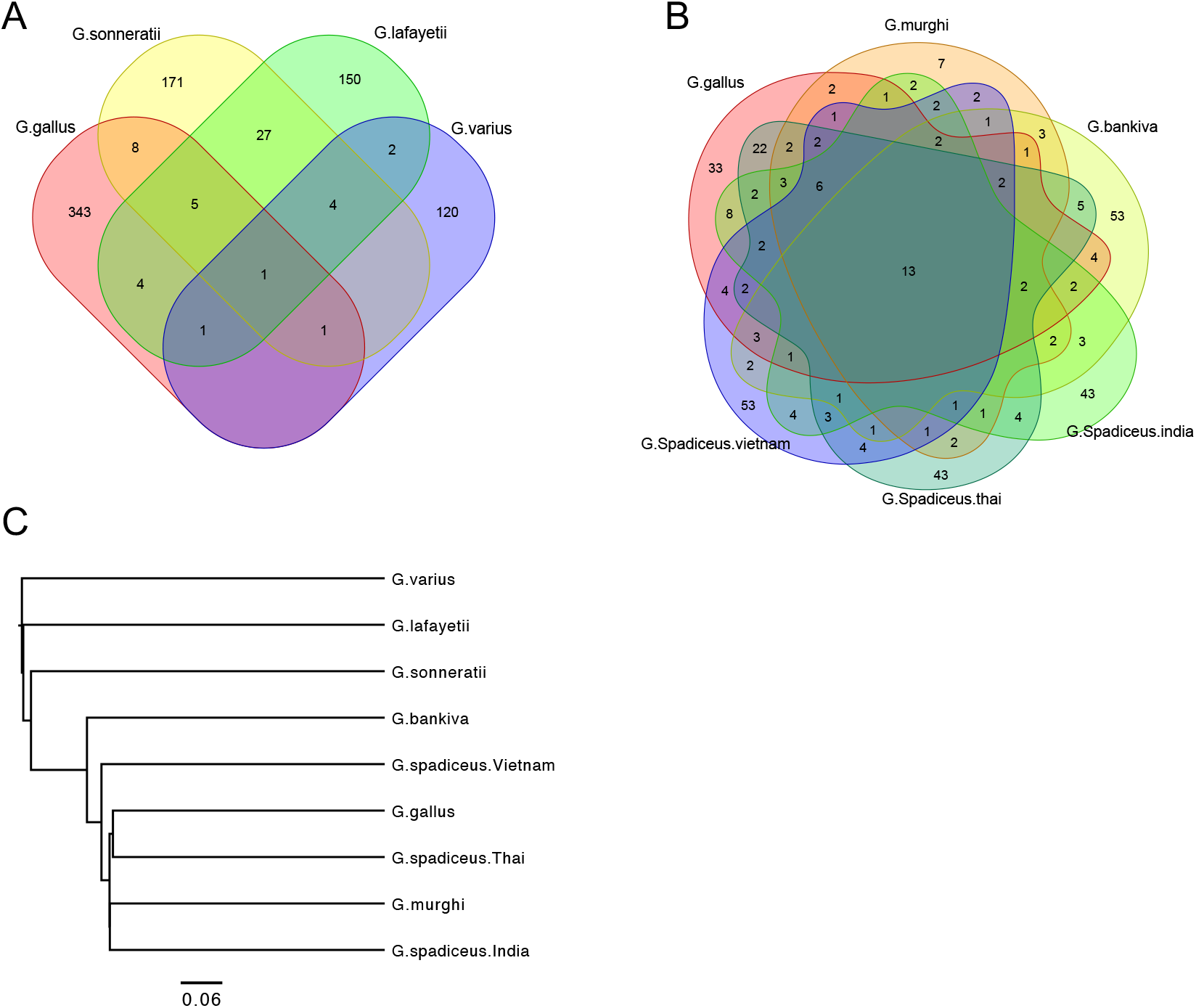
The number of detected endogenous retrovirus (ERV) loci among species and subspecies and the phylogenetic tree. A; Venn diagram indicating the number of ERV loci across four species and the overlap between each ERV loci. B; Venn diagram indicating the number of ERV loci across red junglefowl and its five subspecies and the overlap between each ERV loci. C; A phylogenetic tree constructed based on the presence or absence of ERV loci. The bar indicated each distance.

**Table 1.** The number of Endogenous retrovirus (ERV) loci for each species or subspecies.

| Species/Subspecies | No. of ERV loci |
| --- | --- |
| G.g.gallus | 120 |
| G.g.murghi | 61 |
| G.g.bankiva | 101 |
| G.g.spadiceus.I | 108 |
| G.g.spadiceus.T | 124 |
| G.g.spadiceus.V | 113 |
| G.sonneratii | 217 |
| G.lafayetii | 194 |
| G.varius | 129 |

G.g.Spadiceus.I, G.g.Spadiceus.T and G.g.Spadiceus.V were *Gallus gallus spadiceus* from India, Thai and Vietnam respectively.
The merged number of detected ERV in *Gallus gallus* was 363 loci after removing overlap.

### The classification and the structure of each of the intact ERV-LTRs

A total of 367 loci that were absent in the reference and possessed TSD sequences on both the 5’ and 3’ flanking contig sequences were obtained. Of these, 79 loci were identified across multiple species or subspecies. These loci and its sequences are listed in Table S4. The sequences obtained at the same position were highly similar. For example, among the groups, nine nucleotide substitutions were identified at 346 bp on chr3:99634554. Phylogenetic analysis revealed that 362 of these sequences belong to the LTRs of the EAV family. Simultaneously, the remaining five loci were LTRs of the ALV-E, Ovex1, and MLV-related endogenous retroviruses at the 3, 1, and 1 loci, respectively (Figure 2A). The LTRs of EAV family was further divided into four clusters based on their sequence patterns, with the LTR sequences divided into the U3, R, and U5 regions (Figure 2B and 2C). LTR-D was consistent with EAV-21-3 (Accession No. AJ6232390). LTR-A shared the R and U5 regions with LTR-D and U3 (consistent with U3 of Accession AJ6232391) with LTR-B. LTR-C was consistent with the sequence of M31065 in all regions. Similarly, LTR-B and LTR-C shared identical R and U5 regions, but the U3 regions were distinct.

**Figure 2.**
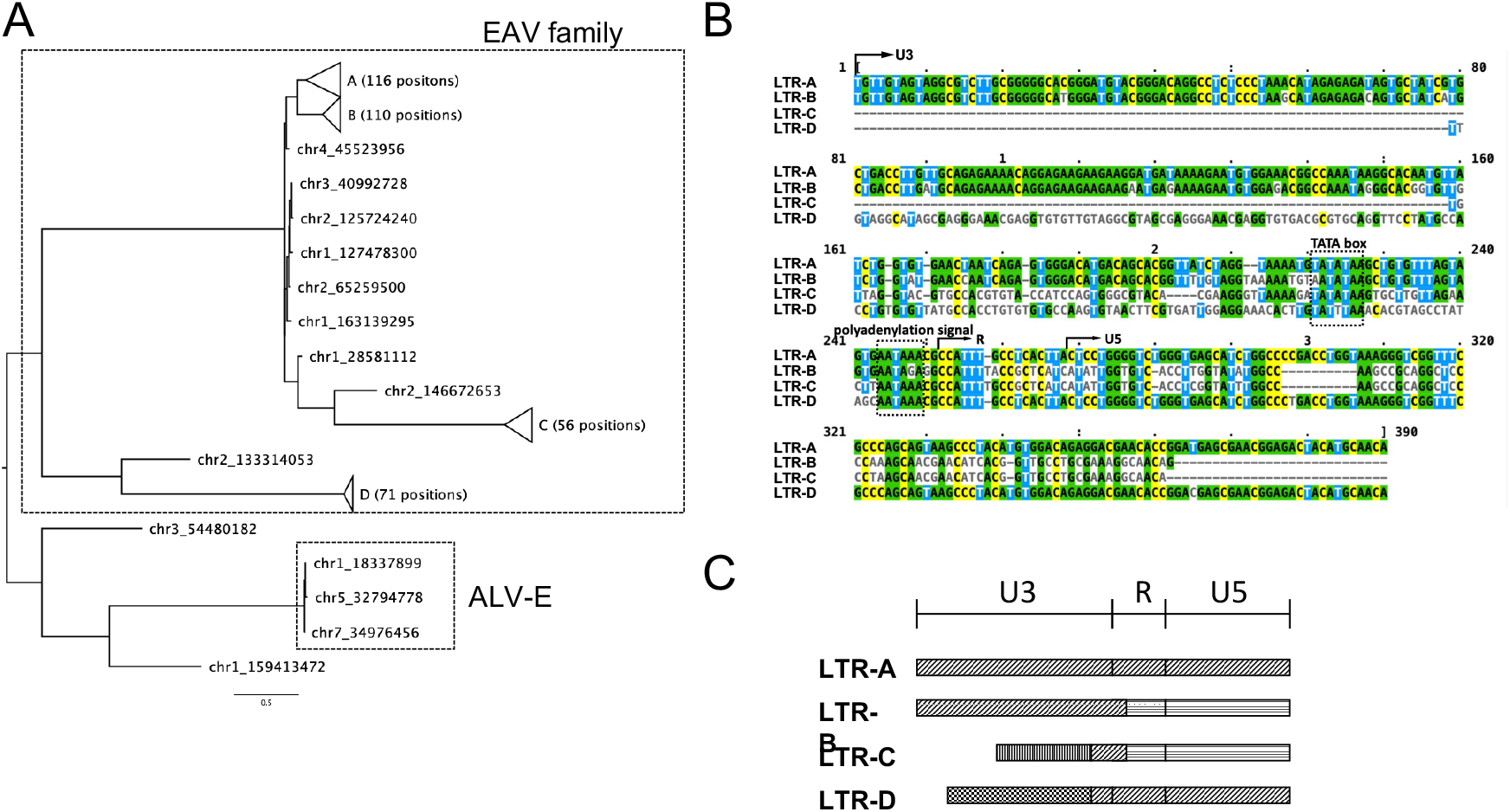
The phylogenetic tree and the structure of each of the intact endogenous retrovirus long terminal repeats (ERV-LTRs). A; A phylogenetic tree constructed based on the long terminal repeat sequence. B; Alignment of the representative sequence from four pattern of Endogenous Avian Virus (EAV) family. C; A schematic diagram of detected EAV-LTR sequence. Identical patterns indicate homologous sequences.

## Discussion

After trimming the sequence data obtained in this study to strict criteria, more than 95% of all read pairs were mapped to the gallus reference genome, although some variation in depth was observed. Thus, the assembled sequence data were considered high quality. In addition, we attempted to detect junglefowl ERVs using improper pairs and singleton sequence reads that did not map correctly to the reference genome. A total of 837 ERV loci were detected in the gallus genome. This result is highly reliable because the presence of TSD was visually confirmed using IGV for the breakpoints detected by RetroSeq, and the contig created by collecting the surrounding sequences also contained ERV-LTR sequences. Previous studies have reported the use of gallus genomes and next-generation sequencing data [20, 21]. For example, one study utilized obsERVer software in conjunction with the Galgal5 reference genome to detect ALV-E in commercial chickens, resulting in the identification of ALV-E at 20 loci [20]. Similarly, 75.22 ± 9.52 integration sites for EAV-HP were identified in commercial chickens, native chickens, and red junglefowl utilized Galgal4 [21]. Although variations in methodologies and reference genomes make direct comparisons difficult, accumulating these findings will undoubtedly contribute to a more comprehensive understanding of the characteristics of endogenous chicken retroviruses. The RetroSeq-based method used in this study primarily targets non-reference ERV loci, which, in theory, are excluded from the reference Gallus gallus genome. As a result, 837 non-reference ERVs were identified at unique genomic positions.

The number of ERV loci identified in the red junglefowl and its subspecies was relatively low compared with the number of ERVs detected in other species. This discrepancy may be attributed to the use of the reference genome of the red junglefowl, which may have resulted in an underestimation of the number of ERVs present, as it does not consider the ERVs unique to the red junglefowl that are already present in the reference genome. Furthermore, the method utilized in this study may have yet to be able to detect all non-reference ERV loci, as a sufficient quantity of improper pairs and singletons is essential for detection. In a specific green junglefowl, a significantly high number of improper pairs (31.59%) was observed. This individual exhibited a higher value (2,011 loci) than other individuals, even after RetroSeq filtering. However, the final ERV loci identified were not significantly different from those of the others, indicating that a certain threshold of data was adequate for detecting non-reference ERVs. Nevertheless, out of 832 locations obtained, only 367 contigs with TSDs on both sides were obtained. This difference may be partly due to the inadequate number of reads. Therefore, increasing the data size may result in the identification of more complete insertion sequences.

The ALV family is younger than the EAV family because it is only found in domestic chickens and red junglefowl, whereas the EAV family is restricted to all Gallus species [22]. This study detected the EAV family in all species, whereas ALV-E was detected only in red junglefowl and its subspecies, which is consistent with previous reports. Therefore, ALV-E is thought to be an internalized sequence in the red junglefowl genome after divergence of the red junglefowl population from the genus Gallus. Species with ERVs at the same locus are thought to have diverged after their common ancestor was infected with a retrovirus and was internalized. A study by [12] estimated the approximate divergence age of the genus Gallus. They calculated that red junglefowl and gray junglefowl diverged 2.56 myr, red junglefowl and Ceylon junglefowl 2.88 myr, gray junglefowl and Ceylon junglefowl 1.77 myr, and green junglefowl and other gallus about 4.0-4.1 myr ago. Overall, the phylogenetic tree constructed from the ERV loci obtained in this study was generally consistent with previously reported phylogenetic relationships. However, it did not reflect the branching age (Figure 1C).

Three loci (chr3:40992728, chr3:101202255, and chr11:7946729) were not consistent with previously reported phylogenetic relationships. For example, on chr3:101202255, ERVs were detected only in red junglefowl, Ceylon junglefowl, and green junglefowl, but not in gray junglefowl. Such ERVs may have been lost from the locus through recombination or other mechanisms during speciation. Alternatively, there may have been instances of introgression between the evolutionarily distant species. Previous research has suggested that introgression from green junglefowl to domestic chickens may have occurred on chromosome 5 [12]. Additionally, whole-genome data analysis has demonstrated an admixture between green junglefowl and red junglefowl species in Indonesia [23].

Comparison of the LTR sequences at the same locus revealed nucleotide substitutions among species and subspecies. Additionally, several sequence patterns were observed in the U3, R, and U5 regions of the LTR of the EAV family in this study. This variation may be a consequence of intra-familial recombination, as previously reported [24]. Although these substitutions and LTR variations do not necessarily reflect genetic divergence, they may support the approximation of the complex history of past introgression. Further examination of the dissemination of ERVs across contiguous regions may enhance our understanding of speciation. In contrast to previous phylogenetic analyses based on sequences mapped to a reference genome, this study employed sequences that do not exist in the reference genome, which may facilitate more detailed phylogenetic analyses in conjunction with previous methods.

In the present study, 306 ERV sequences were detected in the genes, some of which were associated with cell adhesion. The presence of ERVs in the chicken genome affects the host. For example, one of the known effects of ERVs on chickens is the blue eggshell phenotype; the SLCO1B3 gene is expressed in the uterus of hens that lay blue-shelled eggs but not in hens without blue eggshells [8]. An insertion of EAV-HP was identified in the 5’ flanking region of SLCO1B3, and in situ hybridization revealed EAV-HP in the 5’ flanking region of SLCO1B3 [8]. In situ hybridization showed that EAV-HP insertion was associated with the blue eggshell phenotype. In the present study, LTR insertion into cell adhesion-related genes such as RELN, CNTN5, CDH20, CDH7, TENM1, SPON1, NRXN3 and CDH4, was detected. The U3 region of an LTR contains enhancer and promoter sequences that drive viral transcription [25]. It contains other transcription regulatory signals such as the TATA box [26]. The LTR sequence inserted into CNTN5 and NRXN3 contained the TATA box, suggesting that the insertion of ERV LTRs may have played a role in the evolution of cell adhesion processes. Further research is required to completely understand the mechanisms by which ERVs influence the evolution of cell adhesion and other biological processes.

## Materials and Methods

### Whole genome sequence data

Illumina WGS data were obtained in fastq format from the European Nucleotide Archive. Accession IDs are listed in Table S1. Nucleotides with low-quality scores in these reads were trimmed, and adapters were removed with Trimmomatic v.0.36 using the ILLUMINACLIP: TruSeq3-PE:2:30:10, LEADING:3, SLIDINGWINDOW:4:20, and MINLEN:30 settings [27]. The reads were GRCg6a (GCF_000002315.6) using Burrows-Wheeler Aligner and Mem algorithms. Data were produced in the BAM format.

### Detection of non-reference ERVs

ERV detection was performed according to a previous method [15, 16]. We defined the types of read pairs that were mapped to the reference genome and extracted sequence reads that were useful for this study. Most of the paired-end reads were obtained from the WGS map of the reference genome. However, mismatched read pairs can also occur with unexpected span sizes and orientations. Non-proper pairs are those in which the 5’ or 3’ end maps to a contig sequence in the reference genome, and the other end maps entirely or partially to an unexpected locus. Singleton refers to the mapping to the reference genome. A singleton refers to one end of a read pair that does not map to the reference genome, whereas an unmapped read pair refers to both ends of a read pair that does not map to the reference genome (Figure 3). Mismatched read pairs may provide insight into LTR-related loci as anchors. RetroSeq software was used to detect non-reference transposon elements (TEs) using mismatched reads [17]. The process flow is illustrated in Figure 2. The ERV sequences used for RetroSeq were obtained from the National Center for Biotechnology Information (NCBI, Bethesda, MD, USA) and are listed in Table S5. The reference genome was GRCg6a, which contained only autosomes and sex chromosomes. In the RetroSeq “call” step, TE insertion positions (breakpoints) were estimated using reads detected in the “discover” phase as previously reported. The call step was set to ≥10 to reduce false positives, and the maximum read depth option per call was set to 10,000 to increase the BAM coverage. All other retroSeq options were used with their default values. A minimum of seven filter-level breakpoints were used. A breakpoint detected within 500 bp was considered identical and excluded. The IGV was used to detect loci containing TSDs. The loci were presumed to be TSD if they mapped on reads detected during the “discover” phase either from the 5’ or 3’ side, overlapping by 1–10 bp (Figure 3). The 5’ and 3’ reads mapped within 150 bp of TSD were extracted using SAMtools [28]. The extracted read set was used to generate the contig using the CAP3 software [29]. The contig sequences obtained by CAP3 were used for a blast search [19]. The lowest e-value was used to determine ERV class. Each 200 bp sequence upstream and downstream of the breakpoint was extracted from the reference genome and subjected to blastn to eliminate the possibility of detecting ERV sequences in the reference genome. Loci that matched the ERVs were excluded from the analysis.

**Figure 3.**
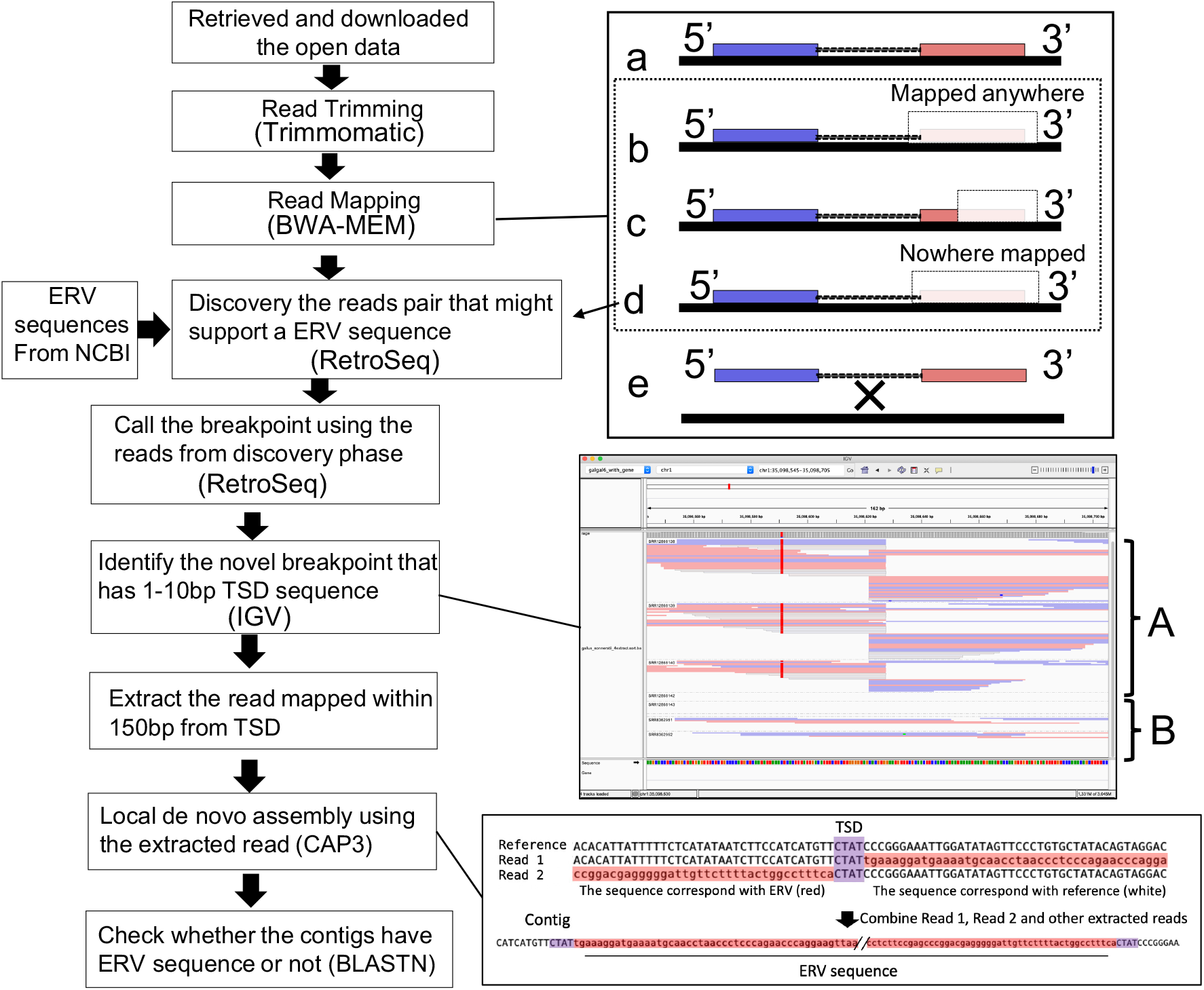
In right-upper panel, black line denotes the chicken reference genome sequence. Blue and red boxes connected with lines denote the 5’ and 3’ ends of a paired-end sequencing read. Most paired-end reads were identified as proper mapping while a small percentage of them were improper mapping. Proper pair; both end of the paired-end sequence mapped accurately (a). Discordant read and split read; one end of the paired-end sequence mapped accurately while the other end was only partially identified at the expected locus on the reference genome. The unidentified sequence could be mapped anywhere else on the reference genome (b and c). Singleton; one end of the paired-end sequence mapped accurately while the other end did not map on the reference genome (d). Unmapped read pairs; neither read mapped to the reference genome (e). Discordant read, split read and singleton were used for RetroSeq analysis. In right-middle panel, a representative view of the integrative genomics viewer (IGV) is used to confirm the presence of target site duplications (TSD) at each locus detected by RetroSeq, extracted support reads from the TSD loci, performed local assembly, and analyzed the contigs for the presence of ERV-genome junctions from both sides. “A” denote the individuals that have the TSD, and “B” denote the individuals that did not have the TSD. The right-lower panel show conceptual diagram of local de novo assembly using cap3. Sequences in red indicate sequences not present on the reference genome, and sequences in purple indicate TSDs.

### Analysis for the obtained ERV

The identified ERV loci were examined for insertion within the gene using IGV. GO analyses for each gene with an ERV sequence were performed using R package clusterProfiler [30]. The author assumed the presence or absence of ERVs at each locus as one or zero for clustering among species and subspecies. The phylogenetic tree of clustering was generated using the function of “dist.binary” by ade4 [30] and “hclust” by ape [32] package of R software [33]. The ERV-LTR sequences of each locus were aligned using ClustalW [34], and a phylogenetic tree was constructed using the maximum likelihood method in MEGA X [35, 36]. Phylogenetic trees and alignments were visualized using FigtTee v1.4.4 (http://tree.bio.ed.ac.uk/software/figtree/) and Mview v1.67 [37], respectively.

## Acknowledgements

This work was supported by Japan Society for the Promotion of Science Grant-in-Aid for Early-Career Scientists Grant Number 22K14907. Computations were partially performed on the NIG supercomputer at ROIS National Institute of Genetics. We would like to thank Editage (www.editage.com) for English language editing.

## Author Contributions

S.I. performed all the experiments, data analysis and writing the final manuscript.

